# Enriching for Extracellular Vesicles from Human Bone

**DOI:** 10.1101/2025.06.18.660234

**Authors:** Christopher J. Wells, Samantha M. Holmes, Evelyn G. Attard, Isabelle Grenier-Pleau, Christine Hall, Eric Bonneil, Pierre Thibault, John F. Rudan, Steve Mann, Sheela A. Abraham

## Abstract

Extracellular vesicles (EVs) are nano-sized membrane-bound structures thought to be secreted by all cells and increasingly recognized as key mediators of intercellular communication. Established EV isolation protocols for bodily fluids-primarily focus on blood with limited insights into methods optimized for EVs from other hematopoietic regions. In this study, we present a novel protocol for the isolation and enrichment of EVs from human trabecular bone and bone marrow. This method employs a two-step purification strategy, combining iodixanol density cushion (IDC) ultracentrifugation with size exclusion chromatography (SEC), and enables EV recovery from fresh tissue hours after collection. Importantly, this approach facilitates the enrichment of bone-derived EVs without the need for enzymatic digestion or long-term culture, preserving native EV populations. This protocol offers a valuable tool for researchers investigating EVs derived from the diverse cellular constituents of the bone microenvironment.

## Introduction

Extracellular vesicles (EVs) are a heterogenous group of nano sized phospholipid particles that are thought to be released by all cells^1–3^. EVs carry bioactive cargo that facilitate communication through molecular signaling and intercellular crosstalk, including proteins, lipids, nucleic acids, and other metabolites^4^. The biological relevance of EVs lies in their ability to transfer molecular information to neighboring cells through their release from donor cells and subsequent uptake into recipient cells, thereby influencing recipient cell function. The role of EVs has been studied in both normal physiological and pathological processes^5–8^. EVs have also been implicated in the bone marrow niche through the regulation of angiogenesis, coagulation, bone calcification, and HSC regulation^9–12^. To date, EVs have been broadly characterized into two main groups based on their biogenesis. Exosomes originate from the endosomal pathway and are released upon the fusion of multivesicular bodies with the plasma membrane, whereas ectosomes are formed through the outward budding and fission of the plasma membrane^13^. Within these broadly characterized groups, there are many EV subpopulations, ranging from 30-1000 nm, including exomeres, small EVs, microvesicles, and apoptotic bodies^14^. Due to the heterogenous nature and overlapping sizes of different EV groups, it can be difficult to accurately distinguish EV populations from each other and from their cell of origin within the microenvironment.

The minimal information for studies of extracellular vesicles 2023 (MISEV2023) provides the most recent guidelines from the International Society for Extracellular Vesicles (ISEV), with standardized framework for the best practice guidelines in EV research^15^. The isolation of EVs can be completed using varying methods, all of which may have differing purities and yields. MISEV2023 guidelines recommend the use of either differential or ultracentrifugation, size exclusion chromatography, and/or density gradients for the purification of EVs. A major hurdle in the enrichment of EVs, especially from blood, is that they are difficult to separate from contaminating lipoproteins and chylomicrons due to their similar phenotype^16^. We implement a 2-step enrichment process consisting of iodixanol density cushion (IDC) ultracentrifugation, in conjunction with size exclusion chromatography (SEC)^16^. This allows particles to be separated first by density and then by size. This combined approach is particularly well-suited for isolating small EVs (<200nm) from plasma, where the extracellular environment is highly complex and rich in both protein and lipid-rich molecules^17,18^. This methodology can also be transferable to other haematopoietic regions, including the bone marrow and trabecular fluid. The use of sequential separation methods enhances the purity of vesicles collected from these tissues, minimizing co-isolation of matrix components, cellular debris, and lipoprotein-like particles. This is critical for accurate downstream characterization and functional studies of small EVs in the bone microenvironment.

### Human Bone Structure

Long bones consist of a combination of dense and porous regions that optimize strength and flexibility within the skeletal system^19^. Bones generally consist of a hard calcified exterior (known as cortical or compact bone) and an inner core of trabecular or cancellous bone. Bones contain marrow along with multiple cell types that include osteoblasts, adipocytes, macrophages, megakaryocytes, regulatory T cells, sympathetic nerve cells, Schwann cells and hematopoietic stem and progenitor cells (HSPCs). Trabecular bone is a porous, lattice-like network found at the epiphyseal and metaphyseal regions that flank the diaphysis^20,21^. In adults, bone marrow can be classified into 2 categories; the yellow marrow consists of fat-rich regions found in the medullary cavity, while red marrow is found in the spongy bone, and is an important site for red blood cell formation^22,23^. The bone marrow relies on sufficient blood supply to maintain adult haematopoiesis. Oxygen and nutrients from circulation can enter the bone marrow through the central artery, which then branches into smaller arterioles^24,25^. Exchange of biomolecules takes place when these arterioles transition into a network of sinusoids and fenestrated vessels throughout the bone space. The central sinus then converges with sinusoids to allow for the release of HSPCs and waste products from the bone marrow into venous circulation^26^. The intricate circulation within the bone marrow and trabecular bone also facilitates the transport of EVs, thereby influencing haematopoiesis, immune regulation, and bone remodelling.

Studying the human bone environment is challenging based on access and rarity of samples. One way of obtaining bone samples is during total hip arthroplasty (THA). This surgical procedure is performed to replace damaged hip joints with a prosthetic implant. In brief, the femoral head is dislocated from the acetabulum and removed using a bone saw. This allows access to the femoral canal (or medullary cavity) where a metal shaft is placed to connect the new prosthetic femoral head^27^. Due to the nature of this procedure, bone marrow aspirates can be retrieved from patients undergoing THA through the medullary cavity. The surgeon may place a syringe into the cavity to collect liquid marrow present throughout the canal. This canal is typically hollowed out to prepare for the placement of the femoral stem, making it part of the routine procedure. The femoral head can also be collected after surgical removal. A bone saw may be used to cut the bone in half, exposing the extensive network of trabeculae within the bone space. This also allows for harvesting of HSPCs and other components specific to the bone microenvironment, such as EVs. This protocol will outline our methodology to enrich for EVs from the bone marrow and trabecular region.

### Protocol Overview

The enrichment of small EVs from bone marrow and trabecular bone is typically completed in tandem. Both isolation techniques use a two-step process: iodixanol density cushion (IDC) ultracentrifugation followed by size exclusion chromatography (SEC) (Fig. 1). Pre-processing procedures differ between bone marrow and trabecular bone due to differences in the biological content, viscosity and state upon collection. Bone marrow obtained from the medullary canal is mostly a viscous liquid, allowing for direct processing through centrifugation and filtration. In contrast, trabecular bone is received as a solid tissue and requires mechanical dissociation to release EVs prior to downstream isolation.

**Figure 1.**
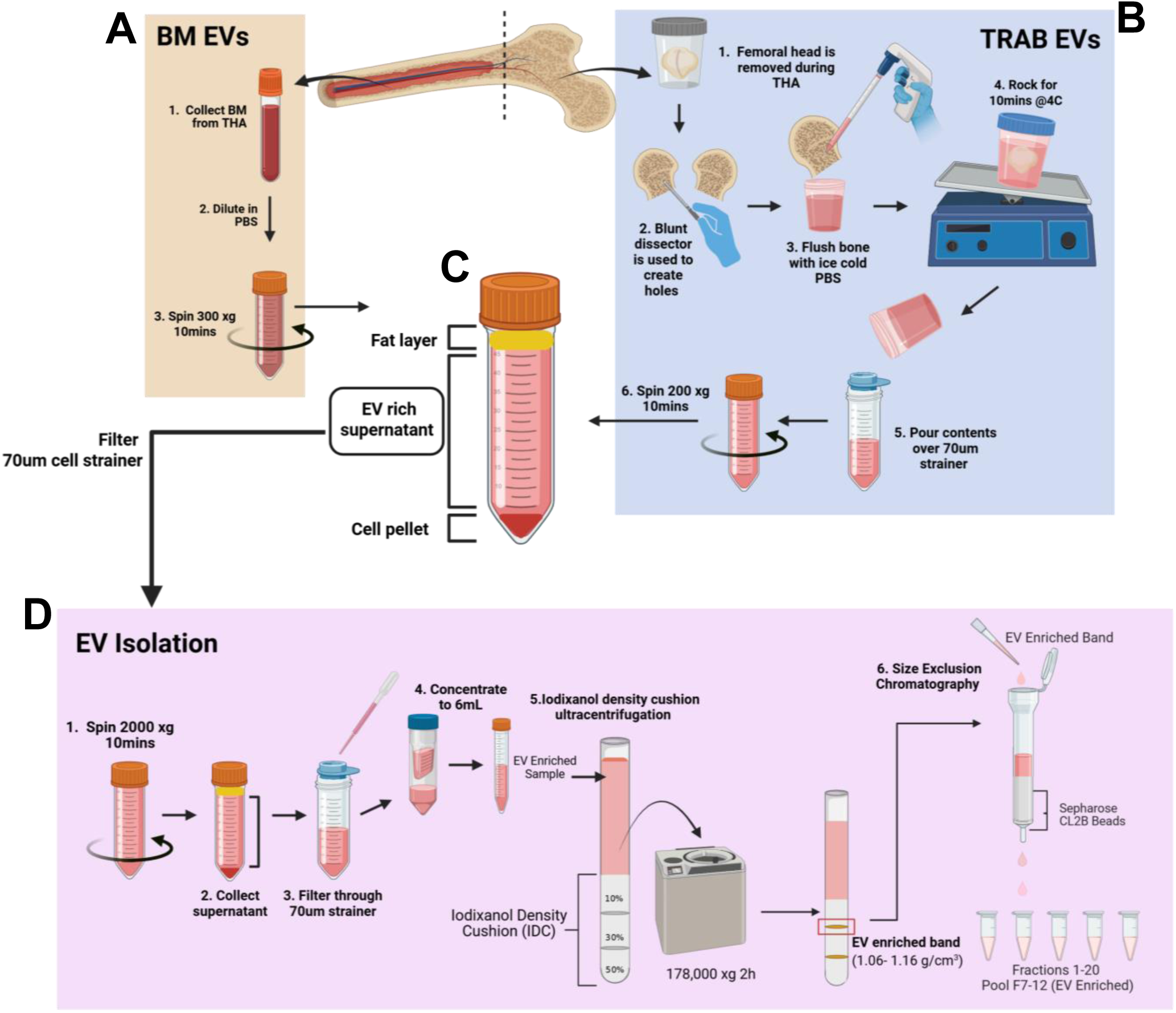
Summary of extracellular vesicle (EV) isolation protocol from bone marrow (BM) and trabecular (TRAB) bone. (**A**) Preparation and isolation of bone marrow biological fluid from total hip arthroplasty (THA). **(B)** Trabeculae preparation for collection of biological fluid via flushing of trabecular bone after boring holes in structure. (**C**) After centrifugation steps for **A** bone marrow and **B** trabecular bone, the EV-rich supernatant is collected and processed prior to two-step EV enrichment. (**D**) Two-step EV enrichment protocols utilizes iodixanol density cushion (IDC) ultracentrifugation followed by size exclusion chromatography (SEC) to separate EVs from contaminating biological fluid components.

After collection, bone marrow samples are diluted in PBS to remove excess fat and centrifuged at 300 x g for 10 min at 21°C (Fig. 1A). The EV-rich supernatant is then collected and washed again prior to EV enrichment. Trabecular bone is probed using a bent-end blunt dissector and flushed with PBS before rocking for 10 min at 4°C to elute remaining EVs within the bone matrix (Fig. 1B). The sample is then strained through a 70μm cell strainer to remove any tissue or debris, and centrifuged at 200 x g for 10 min at 21°C. This cell-free supernatant is then collected and centrifuged at 300 x g for 10 min at 21°C to further remove any remaining contaminants before EV enrichment.

Prior to the 2-step enrichment protocol, samples are first centrifuged at 2500 x g for 10 min to remove any remaining fat, platelets and other debris (Fig. 1D). Bone marrow and trabecular bone samples are then concentrated to a total volume of 6mL using Amicon® Ultra-15 10 kDa filters (Sigma-Aldrich). The 6mL samples are then loaded onto an iodixanol density cushion (50%, 30%, 10%) (OptiPrep, Cedarlane). IDC samples are centrifuged at 178,000 x g for 2 hours at 4°C, using a SW41Ti ultracentrifuge rotor. 1.2 mL is collected from the high-density band located at the interface of the 10-30% iodixanol layers (∼1.06-1.16g/cm^3^) for further processing using SEC columns (CL-2B Sepharose (Sigma-Aldrich)). Samples are collected in 20 0.5mL SEC fractions. Small EV-containing fractions (7-12)^16^ are pooled and aliquoted to allow for further analysis. EV characterization is performed using nanoparticle tracking analysis (NTA) with the goal of evaluating EV size and concentration. Protein concentration can also be determined using the Qubit Protein Assay Kit (Thermo Fisher Scientific). Our laboratory performs both to confirm consistency during enrichment and downstream processing.

### Comparison with Alternative Approaches

Many protocols have been developed for isolating EVs from cell culture supernatants^28,29^ and body fluids, such as blood^16,30–32^ and urine^33,34^, yet relatively few focus on extracting EVs from solid tissues, particularly mineralized tissues like bone. To date, this is the first protocol developed for isolating EVs directly from human bone tissue without the use of cell culture techniques. Traditional approaches for isolating tissue-derived EVs often rely on mechanical homogenization^35,36^ or enzymatic disruption^37^, which can compromise cellular integrity and result in contamination by intracellular vesicles^36^. Our protocol can be used for the enrichment of small EVs (<200nm) from both trabecular bone and bone marrow, without the use of enzymatic treatment or harsh mechanical disruption, therefore limiting the introduction of EV-like artefacts to preserve vesicle integrity and enhance overall purity. To our knowledge, only a limited number of studies have attempted to isolate EVs from bone or bone associated compartments^38,39^, and these typically rely on culture-derived EVs from bone marrow mesenchymal stromal cells^40,41^. Our protocol allows us to process samples immediately after collection, minimizing alterations to the physiological EV composition in different bone regions.

The enrichment of EVs from solid tissues has previously been completed in mostly brain and tumor tissue^37,42–44^. For these methods, enzymatic digestion is typically used, along with tissue homogenization. Enzymatic tissue dissociation – while effective at liberating matrix-bound components – has been shown to induce cellular stress, membrane blebbing, and the release of EV-like artefacts that can confound downstream EV characterization^45^. To retrieve small EVs from the trabecular bone matrices, we use a mechanical flushing technique wherein small perforations are made in the trabecular bone, followed by gentle irrigation with cold PBS, where the eluate is recovered for EV enrichment. This method facilitates the release of interstitial and matrix-associated EVs without altering the physiological vesicle populations. By minimizing ex vivo perturbations, our approach ensures a more physiologically relevant EV profile representative of the in-situ bone microenvironment.

Another key advantage of our protocol is that it enables direct EV enrichment from freshly harvested bone and bone marrow without requiring cell culture or extended incubation of tissue explants. Many existing EV enrichment techniques depend on conditioned media from cultured cell lines or primary cells^46,47^. By processing fresh tissues, we preserve the EV composition and capture vesicles that were actively present within the tissue microenvironment at the time of processing. This strategy not only eliminates artefactual contributions from culture but also ensures that the EV population studies are representative of the true human bone microenvironment and can more effectively replicate their functionality.

Additionally, combining IDC ultracentrifugation with SEC allows for the enrichment of a more comprehensive and physiologically relevant EV profile^15,16^. The two-step isolation procedure allows for the enrichment of small EV isolates from bodily fluids with complex biochemical and physical composition. The use of SEC provides an additional purification step that further enhances purity by removing residual soluble proteins and small non-vesicular particles based on size exclusion principles. Unlike sequential high-speed pelleting, SEC minimizes vesicle damage and loss^48^, thereby preserving the structural and functional integrity of EVs while also incorporating two dimensions of purification: density and size. This protocol – originally adapted for the enrichment of blood plasma EVs – has proven effective not only in various bone regions but also in other haematopoietic-rich sources, such as umbilical cord blood.

### Limitations of Proposed Approach

While the protocol described here offers many advantages for the enrichment of EVs from complex tissue environments, some limitations should be acknowledged. This protocol utilizes one of the most common methods for EV isolation, iodixanol density cushion ultracentirfugtaiton^15^. While this method is effective in reducing contaminant protein co-isolation, it can be time-intensive, requires specialized equipment, and may not fully resolve vesicles with overlapping densities. Similarly, SEC, although a gentler method for EV enrichment, can have limited resolution for distinguishing between similarly sized particles and can result in a more dilute sample, requiring more concentration steps downstream that can lead to additional particle loss. However, when used in tandem, the two methods of separation help refine the purity of the sample being collected, thereby mitigating the limitations of each method alone. This increased purity, however, will likely result in a decrease in overall EV yield after isolation. Sample variability also presents a challenge – differences in vascularization, tissue integrity and cellularity between patients and anatomical regions can impact EV yield. Tissue handling during surgical collection may introduce artefactual EV-like particles due to mechanical disruption. The use of syringes for marrow aspiration or bone cutting during femoral head removal may cause cellular damage and subsequent release of intracellular vesicles, potentially confounding downstream analyses. As this is a routine procedure, all samples collected will be subject to these conditions, potentially mitigating any variation seen. Additionally, the physiological conditions that necessitate surgery (e.g., trauma, malignancy, inflammation) may itself influence EV composition. To minimize the confounding effect from diseased conditions, patients with current or prior diagnosis of haematological disorders and/or active disease are excluded from sample collection.

### Experimental Design

#### Sample Collection – Bone marrow

The protocol described here has been optimized to isolate EVs from both the bone marrow and trabecular bone. Collection of these samples must follow institutional regulations and adhere to the ethics boards associated with human research participants. Due to the nature of these samples, there can be a wide range of variability in the number of EVs present in each sample at the time of collection. For example, bone marrow volumes can range between 5 – 15mL, partly due to the collection of bone marrow as well as the contribution of contaminating wound blood. Bone marrow should be collected in EDTA anticoagulant specimen tubes to prevent clotting during transport (Fig. 2-1). Enrichment of EVs is typically carried out on the day of collection. Alternatively EV enrichment can be performed the following day after being maintained at 4°C, with minimal EV loss.

**Figure 2.**
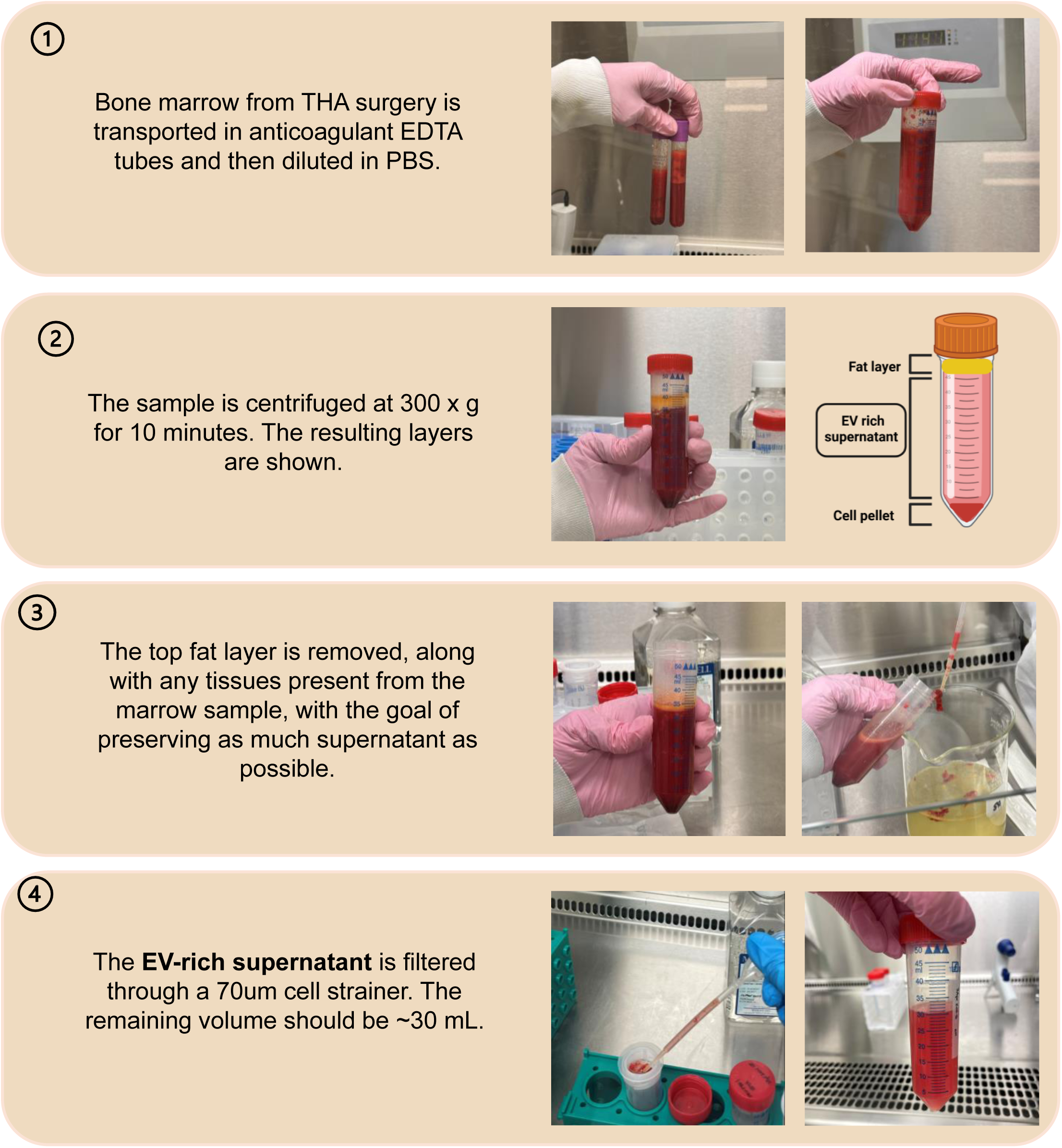
Stepwise protocol for the preparation of EV-rich supernatant from bone marrow.

#### Sample Collection – Trabecular bone

For our protocol, the surgeon cuts the intact femoral head in half for better access to the network of trabeculae within the bone. The bone is placed in a sterile specimen cup for transport back to the laboratory for further processing (Fig. 3-1). The size of the femoral head and volume of accessible bone space varies between male and female patients; however we have not observed sex-specific differences in EV concentration between male and female patients. To improve the retrieval of EVs from the porous bone, we use a bent-end blunt dissector and gently bore holes in the matrices of the distal portion of femoral head (Fig. 3-2). Typically, we have found that the top (proximal) portions of the femoral head are too dense for this type of manipulation and therefore we recommend focusing on the lower (distal) portion near the femoral neck. It is important to be as gentle as possible while probing the trabecular bone to limit intracellular vesicle contamination. Probing of bone is to provide easier access to internal trabeculae and therefore should be kept minimal to avoid disruption.

**Figure 3.**
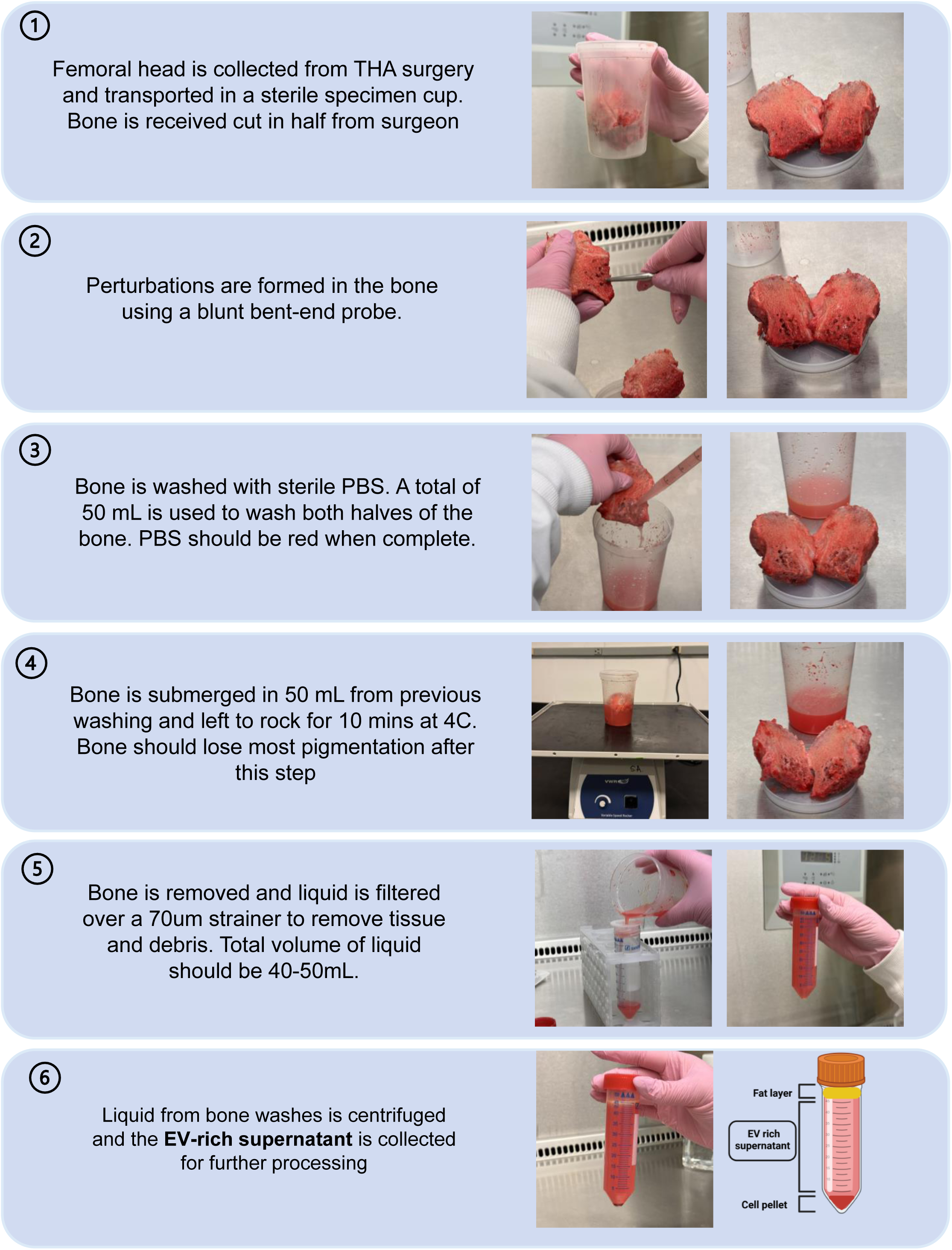
Stepwise protocol for the preparation of EV-rich supernatant from trabecular bone.

#### EV Isolation

The EV-rich supernatants from both bone sources undergo the same enrichment steps after they have been collected (Fig. 4). The volumes (∼45 mL trabecular; ∼30 mL bone marrow) of each supernatant are typically concentrated to a volume of 6 mL before IDC ultracentrifugation to comply with polypropylene centrifuge tubes and SW41Ti rotor specifications of our protocol (Fig. 4-2). For SEC, Sepharose columns should be prepared with degassed PBS a day prior to guarantee proper sedimentation. After isolation is completed, EV-containing fractions 7-12 are pooled and aliquoted in 1.5ml cryovial tubes for storage at −80°C (Fig. 4-5). To avoid multiple freeze-thaw cycles, aliquots can be made. A 250uL aliquot is typically made for each sample for EV characterization, while the rest of the sample is aliquoted for experimental use. Our laboratory uses samples within 6 months of freezing^49^.

**Figure 4.**
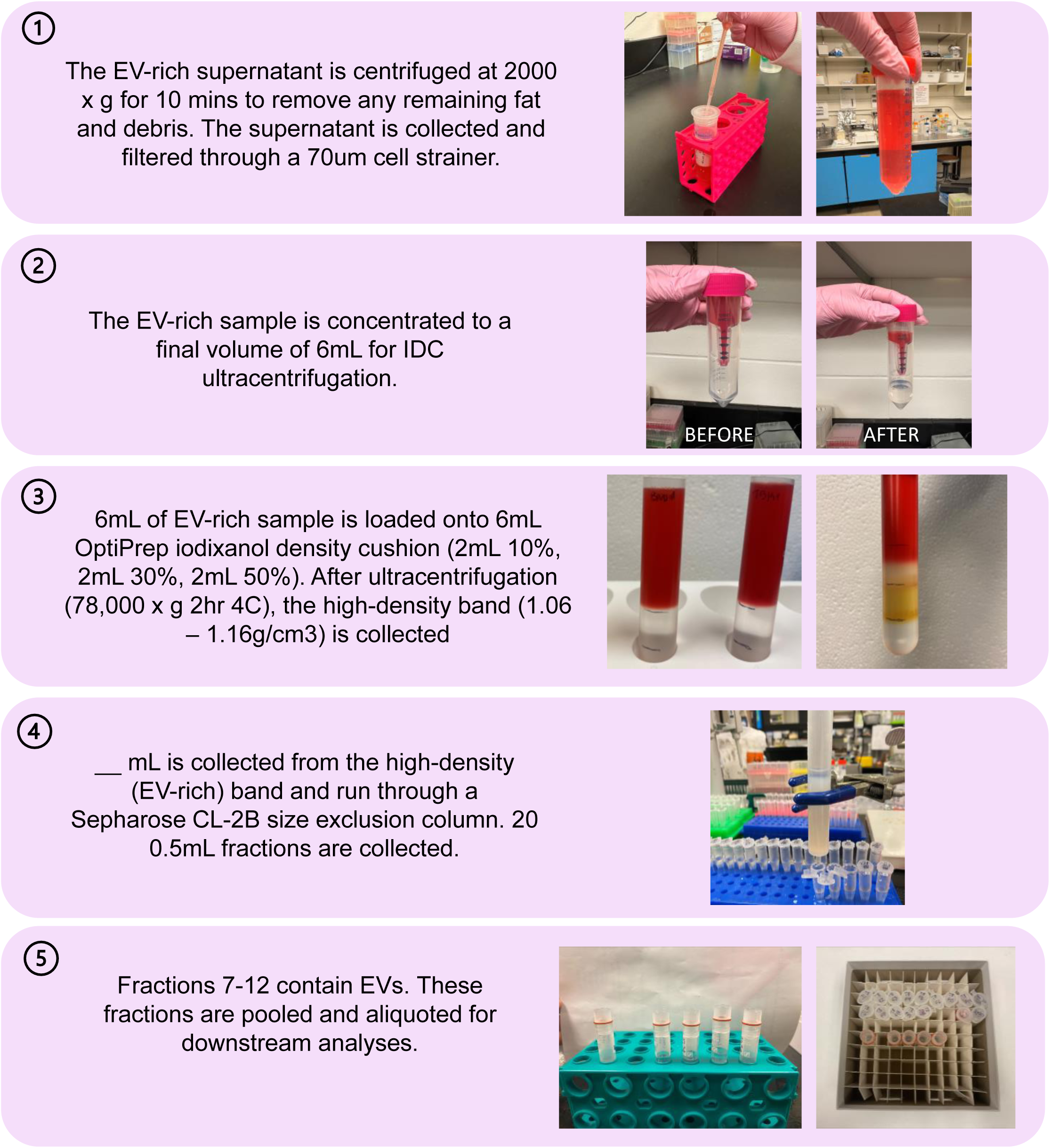
Stepwise protocol for the isolation of extracellular vesicles from bone-related fluids.

**Figure 5.**
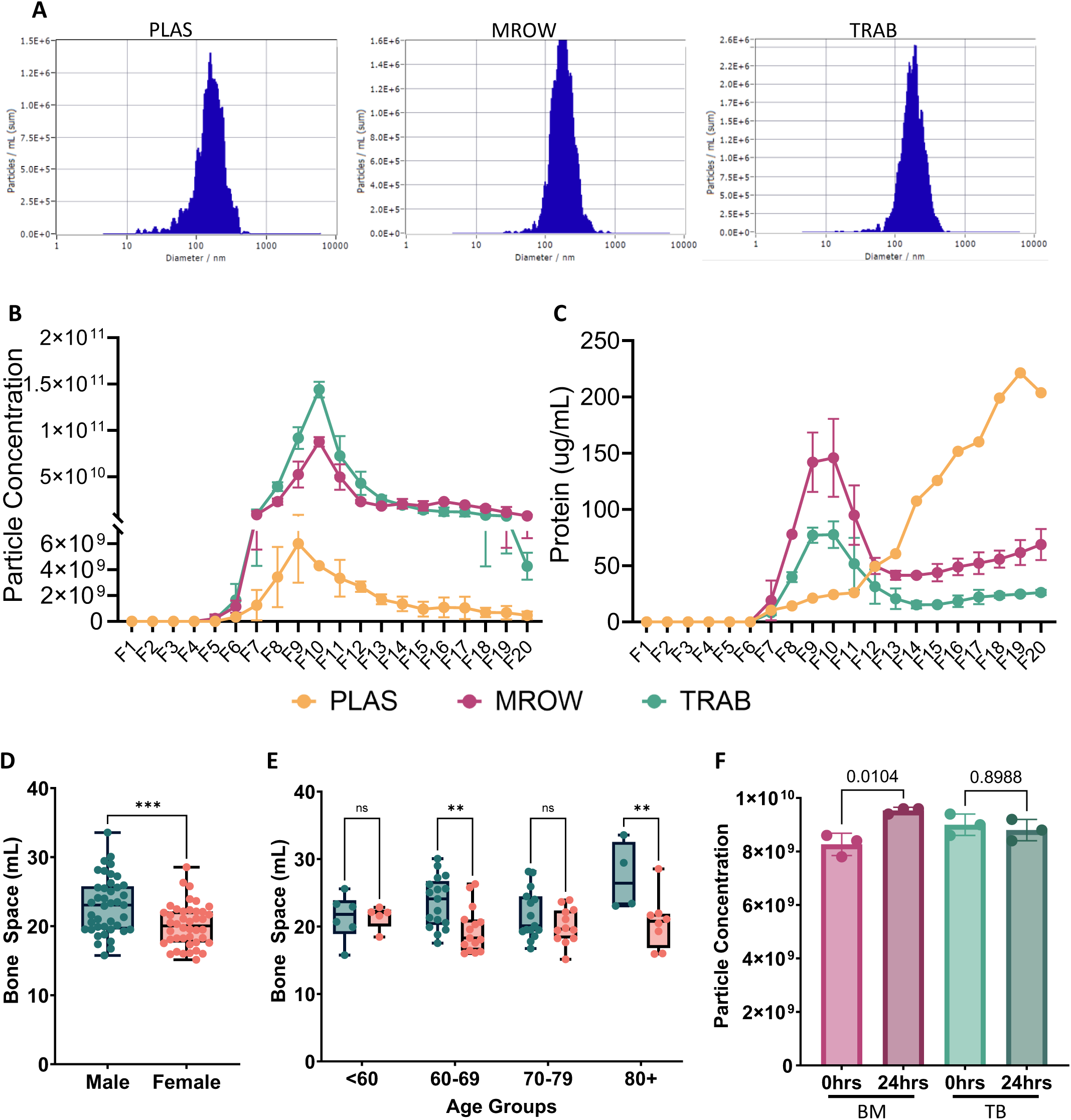
Characterization of EVs isolated from different haematopoietic regions. **(A)** Representative NTA histograms of diameter by concentration (particles per mL). (**B**) SEC fraction analysis particle concentration (n=2). **(B)** SEC fraction analysis by total protein concentration (μg/mL). (**D**) Trabecular bone space comparison between male and female individuals. (**E**) Trabecular bone space comparison separated by age. (**F**) Comparison of EV enrichment from plasma stored for 0hrs and 24hrs at 4°C.

### Biological Materials

#### Human Patients

- Healthy individuals undergoing total hip arthroplasty (THA) surgery. Patients typically range from 43yrs to 85yrs of age. Our human sample collection has followed the Declaration of Helsinki, approved by the Queen’s University Health Sciences and Affiliated Teaching Hospitals Research Ethics Board (HSREB). Informed verbal consent was obtained from all blood donors undergoing total hip arthroplasty surgery according to HSREB regulations.

#### Human Bone Marrow

- The protocol described here has been performed on yellow bone marrow extracted from the proximal end of the femoral diaphysis of individuals undergoing total hip arthroplasty (THA).

#### Human Trabecular Bone

- This protocol has been performed on the red bone marrow eluted from the trabecular bone of femoral epiphysis of patients undergoing total hip arthroplasty (THA).

### Reagents

#### General Laboratory Reagents

- PBS (MultiCell, Wisent Inc., USA, cat. No. 311-010-CL)
- MilliQ distilled H2O (Milli-Q IQ 7000, ThermoFisher Scientific, USA)

#### EV Enrichment

- OptiPrepTM (Serumwek, Germany, cat. No. 1893)
- SepharoseTM CL-2B (Cytiva, Sweden, cat. No. 17014001)
- NanosphereTM Size Standards (ThermoFisher Scientific, USA, cat. No. 3100A)
- Cleaning Solution Concentrate “Green” (Particle Matrix, Germany)

#### Characterization of EV populations

- WB reagents

- RIPA Lysis Buffer, 10X (Millipore, USA, cat. No. 20-188)
- DC Protein Assay Kit II (BioRad, USA, cat. No. 5000112)
- 4X Protein Loading Buffer (LI-COR, USA, cat. No. D01103-01)
- B-mercaptoethanol
- SureCast^TM^ 40% (w/v) Acrylamide (Invitrogen, USA, cat. No. HC2040)
- SureCast^TM^ TEMED (Invitrogen, USA, cat. No. HC2006)
- MES SDS Running Buffer, 20X (Invitrogen, USA, cat. No. NP0002)
- Intercept® Blocking Buffer (LI-COR, USA, cat. No. 927-60001)
- Tween 20
- Exosome-anti-CD63 for Western (ThermoFisher Scientific, USA, cat no. 10628D)
- Anti-CD9 Antibody (Abcam, UK, cat. No. ab92726)
- Anti-Syntenin Antibody (Abcam, UK, cat. No. ab133267)
- Anti-HSP70 Antibody (Abcam, UK, cat. No. ab181606)
- Anti-CD81 Antibody (Abcam, UK, cat. No. ab109201)
- Secondary Antibody (Dylight 800, Dylight 680)
- 1.5M Tris-HCl (pH=8.8)
- 0.5M Tris-HCl (pH=6.8)
- Bis-Tris
- SDS
- Transfer buffer
- Methanol
- ddH2O
- Isopropanol
- 10X TBS
- 1X TBS-T
- 10% APS Solution

### Equipment

#### General Equipment

- Sterile specimen container, 8oz with lid (Avantor, VWR, USA, cat. No. 76532-302)
- Bent-end blunt dissection tool
- K2-EDTA Vacutainer (BD biosciences, UK, cat. No. 366643)
- 70μM sterile cell strainer (Fisher Scientific, USA, cat no. 22363548)
- Polypropylene tubes (50ml, Sarstedt, Germany, cat. No. 62.547.205)
- Polypropylene tubes (15mL, Sarstedt, Germany, cat no. 62.554.205)
- Microcentrifuge tubes (1.5mL with cap, polypropylene)
- Transfer pipette (Fisher Scientific, USA)
- CryoPure Tube (1.6mL; Sarstedt, Germany, cat no. 72.380)
- Microbiological safety cabinet class II
- Variable Speed Rocker (VWR, USA, BR2000-GM)
- Cold room
- Centrifuge

- Un-refrigerated Heraeus Megafuge 16, with Rotor TX-400 (ThermoScientific, Germany, cat. No. 75004231)
- Refrigerated Heraeus Megafuge 16R, with Rotor TX-400 (ThermoScientific, Germany, cat. No. 75004271)
- P1000 pipette + tips
- Serological pipette
- Tube rack

#### EV Enrichment

- Amicon® Ultra-15 Centrifugal Filters (Millipore, Ireland, cat. No. UFC901024)
- Polypropylene Centrifuge Tubes (14mm x 89mm, Beckman Coulter, USA, cat. No. 331372)
- LS-60M Ultracentrifuge (Beckman coulter, USA, cat. No. 347240)
- SW 41 Ti swinging-bucket rotor (Beckman coulter, USA, cat. No. 331362)
- Econo-Pac Disposable Chromatography Columns (10mL; BioRad, USA, cat. No. 7321010)
- 50mL beakers

#### Particle Measurement

- ZetaView PMX-120 NTA (Particle Matrix, Germany)

#### Protein Measurement

- Qubit 4 fluorometer (ThemoFisher Scientific, USA, cat. No. Q33238)
- Qubit assay tubes (ThemoFisher Scientific, USA, cat. No. Q32856)

#### Western Blotting/Flow

- Microtest plate 96 well flat (Sarstedt, Germany, cat. No. 82.1581)
- Butane Torch
- Absorbance reader
- ThermoMixer C (Eppendorf, Germany, cat. No. 5382)
- Mini Gel Tank (Invitrogen, USA, cat. No. A25977)
- PowerEase 300W (Life Technologies, cat. No. PS0300)
- Mini Blot Module (Invitrogen, USA, cat. No. B1000)
- Immuno-Blot® PVDF Membranes (BioRad, USA, cat. No. 1620177)

## Equipment Setup

### ZetaView PMX-120 NTA

- Syringe 30mL air into cell assembly to remove excess liquid. Remove cell assembly and unscrew cell carrier. Slide out cell carrier and begin cleaning protocol (as outlined by Particle Matrix). Clean the interior of the cell with washing solution (ddH2O, cleaning agent) and pipe cleaner. Rinse with ddH2O before drying with lint-free wipe and compressed air. Once dry, wipe lenses with 95% EtOH and cotton swab. Return cell carrier to cell assembly and screw into place. Reattach cell assembly and follow NTA startup procedures as outlined by ZetaView software.

### Sample Collection

#### Surgical Removal (Conducted by Medical Staff) – Timing variable (b/w 30-75mins)

1. Femoral head is removed, and an anterior cut is made along the femoral head using an oscillating bone saw, to allow access to inner trabeculae.
2. Trabeculae halves are placed into an 8oz sterile container and passed to the researcher.
3. A modified T-handle femoral canal finder rasp is inserted into the bone marrow cavity, while a 50mL syringe extracts any exiting marrow.
4. Remove rasp and insert syringe into cavity for collection of the bone marrow sample.
5. Bone marrow syringe is capped by medical staff and handed to the researcher. *\*Sample collection volume ranges between 2-16mL, depending on biological variability of samples and contaminating wound blood\**

#### Sample Collection and Transport – Timing ∼ 5 mins

6. Uncap the bone marrow syringe, and transfer the sample into K2-EDTA (K2E) tubes
7. Rinse syringe of remaining bone marrow sample with 5mL of PBS and transfer to K2E tubes and invert.
8. Place bone marrow and trabeculae samples into biohazard transport bags before returning to biological safety cabinets.

#### <u>Sample Washing</u> – Timing 60mins

##### Bone Marrow Preparation (Fig. 2)

9. Transfer bone marrow from K2E tubes to 50mL conical tube. Determine and log bone marrow volume (accounting for 5mL PBS during collection).
10. Wash K2E tubes with PBS and transfer to 50mL conical tube.
11. Dilute bone marrow in 40mL of PBS and centrifuge sample at 300 x g for 10mins at 21°C.
12. Remove and discard the top fat layer. Gently pass bone marrow fluid (red middle layer) through a 70μm cell strainer into a new 50mL conical tube. \**If any chunks remain in fluid layer, remove and discard\**
13. Proceed to Fluid Concentration steps

##### Trabecular Fluid Preparation (Fig. 3)

14. Open trabeculae container and add 25mL of PBS.
15. Using a bent-end blunt dissector, bore holes into one half of the trabeculae of the femoral head. Be careful not to break the bone by over-boring. Holes are created to provide easier access to deeper trabeculae without causing mass cell apoptosis. *\*Areas closer to metaphysis of femur are more accessible than the epiphysis\**
16. Rinse the trabeculae ∼20 times with PBS using a serological pipette (10mL/25mL). *\*Should see visible colour change of trabecular bone and wash supernatant from clear to bright peach/red\**
17. Repeat steps 15-16 with the other half of trabeculae.
18. Once washing is complete, place both halves of the trabeculae into the container and cover with PBS (30-50mL).
19. Place container on rocker for 10mins at 4°C.
20. Remove trabecular halves and pour supernatant through 70μm cell strainer into 50mL conical tube. Centrifuge at 200 x g for 10mins at 21°C. *\*Depending on size of femoral head, two conical tubes may be required for the amount of PBS used. Two strainers may also be required depending on the fat and debris of sample\**
21. Remove and discard top fat layer. Transfer trabecular fluid into a new 50mL conical tube.
22. Proceed to Fluid Concentration steps

#### <u>Fluid Concentration</u> – Timing variable (1-2hrs depending on sample) (Fig. 4)

*\*Recommended to keep samples on ice or at 4°C after isolation of fluid*

23. Centrifuge bone marrow and trabecular fluid at 2000 x g for 10mins at 21°C.
24. Remove and discard excess fat layer and transfer fluid into new 50ml conical tubes. Avoid disturbance of remaining cell pellet.
25. Prepare Amicon Ultra-15 centrifuge tubes: Add 10mL ddH_2_O above the filter and centrifuge at 2000 x g for 10mins at 21°C. Remove liquid and repeat the wash step with 10mL PBS.
26. Add 15mL of biological fluid above the centrifuge filter and centrifuge at 3000 x g at 21°C. Centrifugation time is dependent on the amount of sample needed to concentrate. On average it takes ∼30mins to concentrate 10mL of sample. After centrifugation, remove eluted supernatant (keep supernatant above filter) and top up with remaining biological fluid. Repeat centrifugation until sample is concentrated down to final volume of 6mL.

*\*Concentrating time will vary on sample size and between biological replicates\**

#### <u>Preparation of Size Exclusion Columns</u> – Timing ∼ 40mins (suggest preparing day prior)

27. Prepare 200mL of degassed PBS
28. In the BSC, add 15mL per column of Sepharose^TM^ CL-2B to a glass flask and degas using the vacuum.
29. Load 15mL of Sepharose into chromatography column. Wait 20-30mins to sediment.
30. Wash column with 10mL degassed PBS and elute through column. As PBS level gets close to the top of Sepharose, repeat wash step for a total of 3 washes. *\*Do not allow Sepharose^TM^ to dry. If drying occurs, columns should not be used\**
31. After wash steps, plug the column and cap until sample is prepared. *\*If Sepharose is uneven, you can use a transfer pipette to level\**

#### Extracellular Vesicle Enrichment – Timing ∼ 5hrs

32. Prepare iodixanol density cushion by diluting OptiPrep in ddH_2_O to 50%, 30%, and 10% dilutions.
33. Layer 6mL of bone marrow or trabecular sample on top of 2mL 50%, 2mL 30%, and 2mL 10% OptiPrep cushion into a polypropylene ultracentrifuge tube. Make a mark between each layer for better visibility after centrifugation steps.
34. Place sample tubes into SW41Ti swinging buckets and ensure buckets are balanced.
35. Centrifuge samples at 178,000 x g for 2hrs at 4°C (38,000RPM, 2hrs, 4°C on SW41Ti in LS-60M Ultracentrifuge, Beckman Coulter, USA)
36. After spin, there should be 3 visible bands between each layer. Collect 1.2mL of high-density band (HDB) located between the 10-30% layers. This band is enriched for EVs and high-density lipoproteins between 1.06-1.16g/cm^3^. *\*Depending on sample, HDB may be hard to see\**
37. Load the HDB onto the size exclusion chromatography column and collect 20 fractions of 0.5mL into microcentrifuge tubes. Continue to rinse with cold degassed PBS (3mL at a time) until 20 fractions have been collected. *\*Do not allow column to run dry. If drying occurs, columns should not be used \**
38. Fractions 7-12 are EV-containing fractions and should be pooled before EV characterization.

#### Storage of Extracellular Vesicles

39. Aliquot pooled fractions 7-12 into 1.6mL cryovials and store at −80°C. *\*EVs are susceptible to degradation/functional changes upon thawing. Try to minimize freeze thaw cycles by aliquoting appropriately and including aliquot for measuring EV and protein concentration. EVs demonstrate reduced functionality after 6 months of storage at −80°C and therefore, should be used within this timeframe*^49^*.\**

#### Calculation of volume

50. To calculate the volume of the femoral sample, the femoral sample is weighed after sample washing and collection of trabecular plasma. Briefly, after sample collection, trabeculae is dried for 7-10 days. Dry trabeculae is then re-weighed, followed by a rehydration by total submersion in ddH_2_O for 24hrs. Mass of rehydrated trabeculae was determined before disinfection and removal of bone. Calculation of available space within the trabecular bone structure was performed as follows:

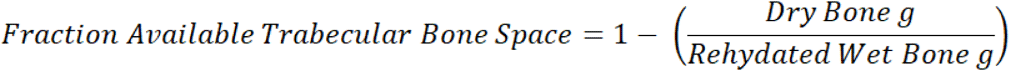

We have calculated the average volumes for males and females separately, generating sex-specific “Fraction Available Trabecular Bone Space” factor that we use in trabecular bone space volume calculations.

To calculate the volume of space available within the trabecular head that was used to recover extracellular vesicles, we use the following calculation:

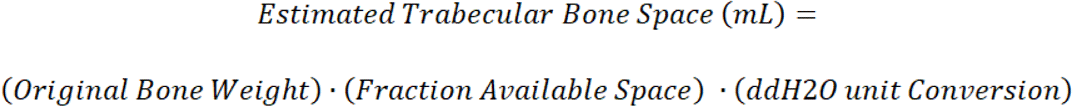

Where fraction available trabecular bone space for females = 0.258, and for males = 0.203. The ddH_2_O unit conversion is a constant at 1.002 mL/g. Example: for a female with a trabecular weight of 71 g, the estimated trabecular bone space calculation would be: (71 g) ’ (0.258) ’ (1.002 mL/g) = 18.35 mL.

#### Anticipated Results

In this protocol, we identify a novel method of isolating EVs from two locations within primary bone tissue. This method highlights the preparation of bone marrow and trabecular biological fluid prior to EV enrichment with a two-step IDC/SEC protocol. After enrichment, EV samples can be characterized. To verify separation of EVs by size, nanoparticle tracking analysis can be conducted across all 20 0.5mL SEC fractions (Fig 5A). This demonstrates that this method for enrichment of EVs consistently elutes between fractions ***<u>F7-12</u>*** for all locations and effectively excludes contaminating soluble proteins from bone fluid (Fig. 5B-C). Characterization of these samples emphasizes the heterogeneity of EV composition across different haematopoietic regions.

The bone space was calculated to determine the volume of trabecular EV samples. Difference in bone space calculations were then compared between sexes and across different age groups. Sex differences were observed with male samples demonstrating significantly larger overall bone spaces in trabecular bone than females (Fig. 5D). These differences are driven by significant differences in male and female bone space volumes within 60-69yr olds and 80+yr old individuals (Fig. 5E).

Proteomic profiling analysis on EVs can be conducted. We have probed for classical EV markers as outlined by online EV databases (EVpedia/ExoCarta/Vesiclepedia) and demonstrate that bone-derived EVs isolated with this method are positive for key tetraspanins (CD9, CD63, CD81, CD47, TSG101) used to confirm EV enrichment in plasma samples (Fig. 6A). To further validate the positive enrichment of EVs from bone, Western blot analysis can be performed for foundational EV markers (CD63, CD9, CD81, HSP70, Syntenin) (Fig. 6B-F). We observe key tetraspanins expressed in EVs from all 3 haematopoietic regions, with bone EVs expressing higher levels of CD63 (Fig. 6B), CD81 (Fig. 6C), and CD9 (Fig. 6D), consistent with proteomic data. Similar levels of expression were observed across haematopoietic regions for the cytosolic proteins HSP70 (Fig. 6E) and Syntenin (SDCBP) (Fig. 6F) were also noted.

**Figure 6.**
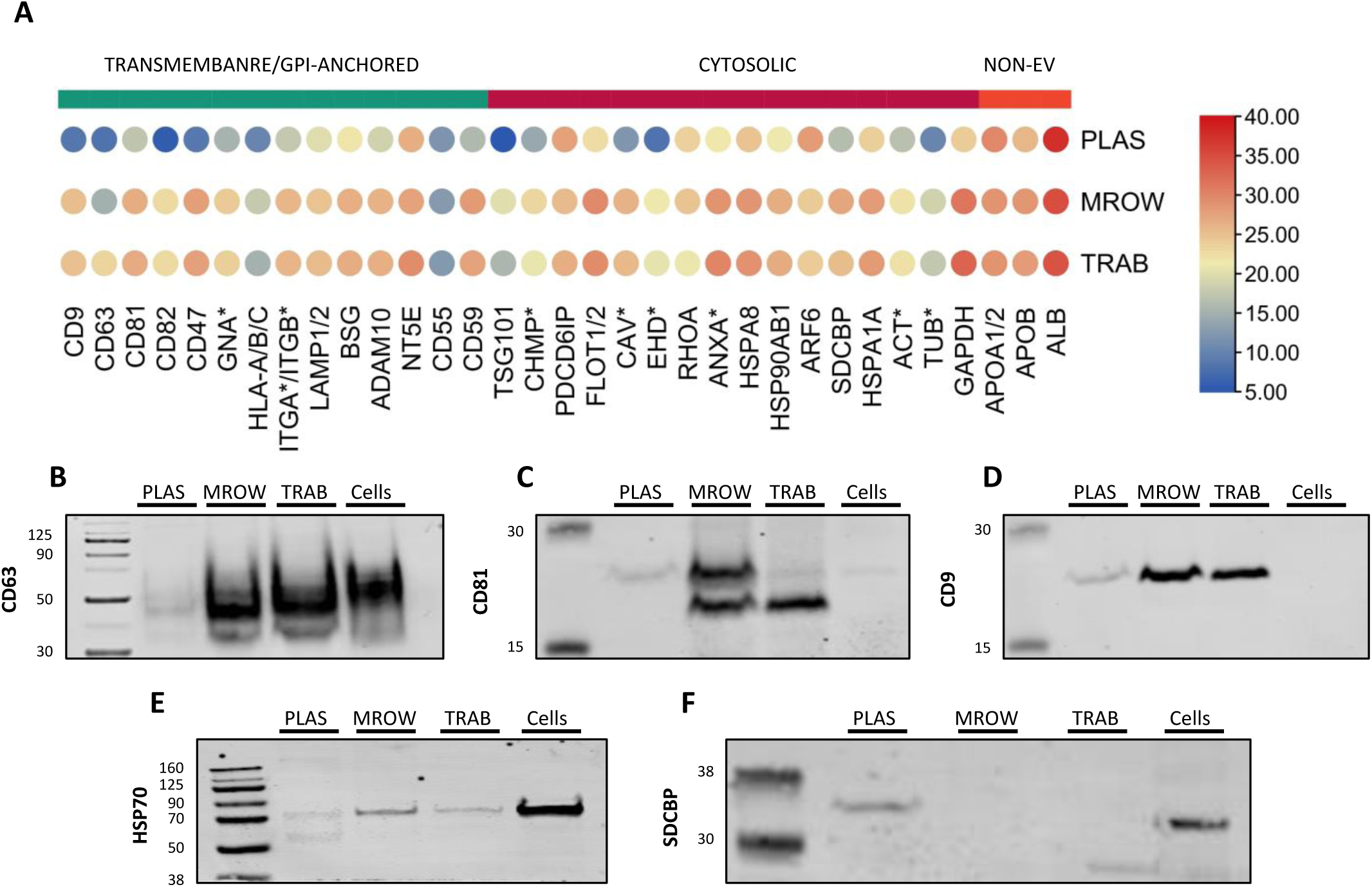
Protein composition and validation of extracellular vesicle populations. Analysis of key EV identifying markers. (**A**) Protein expression data of key EV markers pulled from online databases. Western blot validation of key tetraspanins (**B**) CD63, (**C**) CD81, (**D**) CD9 and cytosolic EV markers (**E**) HSP70, and (**F**) SDCBP (Syntenin).

##### Notes

1. During washing of the bone, supernatant will shift in colour towards a bright pink/red while the bone begins to pale, demonstrating proper elution of trabecular bone marrow. (Fig. 3-4).
2. To provide flexibility in the protocol timeline, we assessed the impact of storing overnight at 4°C. After storage of bone marrow and trabecular fluid at 4°C for 24hrs (up to step 22), two-step EV enrichment was conducted to compare the effect of storage on particle concentration. We have observed a difference in EV concentration within the marrow samples but not the trabecula after being refrigerated (4°C) overnight (Fig. 5F). No differences in size were observed in either marrow or trabecula after overnight refrigeration.
3. It should be noted that we have not tested the effects of freezing the EV-rich supernatant on particle number or size. During processing, samples were always kept on ice where possible.

